# Identification of the Down syndrome critical region 3 gene as a mammalian cell size regulator

**DOI:** 10.64898/2026.07.23.740440

**Authors:** Kazumi Kimura, Masakazu Souda, Ryotaro Mori, Yoshimi Kato, Hiroki Kurahashi, Masashi Asai, Kazuo Yamamoto

**Author notes:** **Correspondence:** Kazuo Yamamoto, Masashi Asai.

## Abstract

Using a genetic screening approach based on an inducible gene-activating system and cell sorting, Down syndrome critical region 3 (DSCR3) was isolated as a gene whose overexpression increased cell size. Fibroblasts derived from individuals with Down syndrome (DS) exhibit elevated DSCR3 expression at both the mRNA and protein levels, correlating with increased cell volume compared to fibroblasts from healthy donors. Despite a slower proliferation rate, DS fibroblasts demonstrate higher basal and maximal mitochondrial respiration, suggesting enhanced metabolic activity associated with increased cell size. siRNA-mediated knockdown of DSCR3 reduces cell size in both DS and normal fibroblasts, indicating its general role in cell size regulation. As DSCR3 is a component of the retriever complex involved in endosomal cargo recycling, these findings position membrane protein trafficking as a novel module for cell size control.

## Introduction

Mammalian cells regulate their size to maintain proper cellular function and overall organismal health (Chadha et al., 2024). Cell size is correlated with intracellular biomolecular scaling, organelle homeostasis, and cell cycle progression. It also affects early development and cell differentiation (Chadha et al., 2024; Li et al. 2015). The optimal cell size is critical for maximizing the growth rate and mitochondrial metabolism, with intermediate-sized cells within a population exhibiting the highest metabolic activity (Miettinen et al. 2017). However, aberrant increases in cell size, such as those caused by polyploidy, can be disadvantageous to cellular metabolism, fitness, and functionality, suggesting that cell size control may have evolved as a guardian of cellular fitness and metabolic activity in eukaryotes. Indeed, cellular hypertrophy can potentially predispose individuals to or worsen metabolic diseases (Miettinen et al., 2017). The importance of cell size control is evident across various organisms, from bacteria to mammals, with different mechanisms employed to maintain size homeostasis (Chadha et al., 2024; Proulx-Giraldeau et al., 2022). Therefore, understanding these mechanisms has significant implications for both diagnosis and treatment in clinical settings.

Several genes have been identified to play important roles in regulating mammalian cell size. For example, overexpression of the tumor suppressor genes TSC1 and TSC2 triggers cell size reduction, whereas a dominant-negative TSC2 mutant induces an increase in cell size (Rosner et al., 2003) as a consequence of its role in the mTOR pathway. Inhibition of mTOR complex 1 components reduces cell and organ sizes (Kim et al., 2002; Porstmann et al., 2009; Sengupta et al., 2010). The transcription factor YAP in the Hippo signaling pathway regulates cell size and growth dynamics (Mugahid et al., 2020; Pérez-González et al., 2019). In addition to these cellular signaling pathways, c-Myc transcription factors have been shown to play an important role in mammalian cell size regulation in the context of tumorigenesis (Rosner et al., 2003). Several models have been proposed to explain cell size regulation in yeast, plants, and mammalian cells (Xie et al., 2022); however, additional research is required to fully understand cell size regulation.

Therefore, we performed genetic screening to identify direct cell size regulators (Yamamoto et al., 2014; Yamamoto & Mak, 2017). One candidate gene, Largen, regulates cell size by enhancing the mitochondrial activity (Yamamoto et al. 2014). In addition to Largen, several other candidates have been confirmed to directly regulate cell size, as their overexpression induces an increase in cell size. Here, we describe the Down syndrome critical region 3 (DSCR3) gene as a cell size regulator that contributes to the cell size of individuals with Down syndrome (DS).

## Results

### Identification of DSCR3 as a cell size controlling gene via the genetic screening

We previously reported a genetic screening system to identify possible cell size-controlling genes by combining an induced genomic gene-activating system with cell sorting (Yamamoto et al., 2014, 2017). We established a cell line, JET7, stably expressing the tetracycline-responsive transactivator tTA from the human lymphoma cell line Jurkat and applied it to the infection of a recombinant retrovirus in which the genome contained a small element, ERM-tag, consisting of the HA-epitope tag followed by a consensus splice donor sequence under the control of the tetracycline (Tet)-responsive promoter (Liu et al., 2000). Once the retroviral genome was inserted, the ERM-tag was transcribed from the Tet-responsive promoter by tTA and fused to an exon of the downstream endogenous gene owing to the presence of the splice donor sequence. Thus, we obtained a mutant cell pool in which endogenous genes in the genome were activated in a tTA-dependent manner. The mutant cell pool was then treated with rapamycin. Rapamycin is a selective inhibitor of the growth-directed protein kinase mTOR and has been shown to reduce cell size when added to the culture medium. We hypothesized that if the activation of a particular gene by the ERM retrovirus was sufficient to overcome the cell size reduction induced by rapamycin, the cells containing this gene would remain larger than the other cells. These mutant cells can be collected by sorting large cells. Single-cell clones were established by culturing the sorted cells at a limited dilution, and pseudo-positive cells were excluded by the addition of doxycycline, a stable derivative of Tet, to individual cultures to determine whether the large cell phenotype was reversed by the inactivation of tTA. The fusion transcripts in the positive cell clones were sequenced using reverse transcription (RT)-PCR and 3’RACE to identify the ERM-targeted genes.

Using this strategy, we isolated a few hundred cell clones and identified the possible responsible genes in individual cell clones (Yamamoto et al., 2014). In this study, we characterized one of the candidate cell size-controlling genes identified in the clone 1C6. As shown in Figure 1A, the size of the parental JET7 cells (distribution represented by a black line in the left panel) was reduced in the presence of rapamycin (blue line, left panel). The cell size distribution of clone 1C6 slightly shifted to a larger area (compared to the mean FSC-A values between JET7 and 1C6). Rapamycin also reduced the cell size of clone 1C6; however, its distribution was still larger than that of JET7 (blue line and mean FSC-A value shown in the right panel). The addition of doxycycline to the culture medium of JET7 had no effect on cell size distribution (red line, left panel), even in the presence of rapamycin (green line, left panel). However, the size distribution was slightly shifted to a smaller area in the case of 1C6 cells treated with doxycycline (red line, right panel) and further reduced in the presence of rapamycin (green line, right panel). JET7 and 1C6 cells were transiently transfected with a plasmid expressing mCherry-Cdt1 to label the G1-cell populations. As shown in Supplementary figure 1B, the cell size distribution of G1-gated 1C6 was larger than that of G1-gated JET7, and this trend was almost the same when cell size distributions were compared with non-gated cells. Thus, the cell size of clone 1C6 was controlled by the expression of the gene tagged with ERM.

**FIGURE 1.**
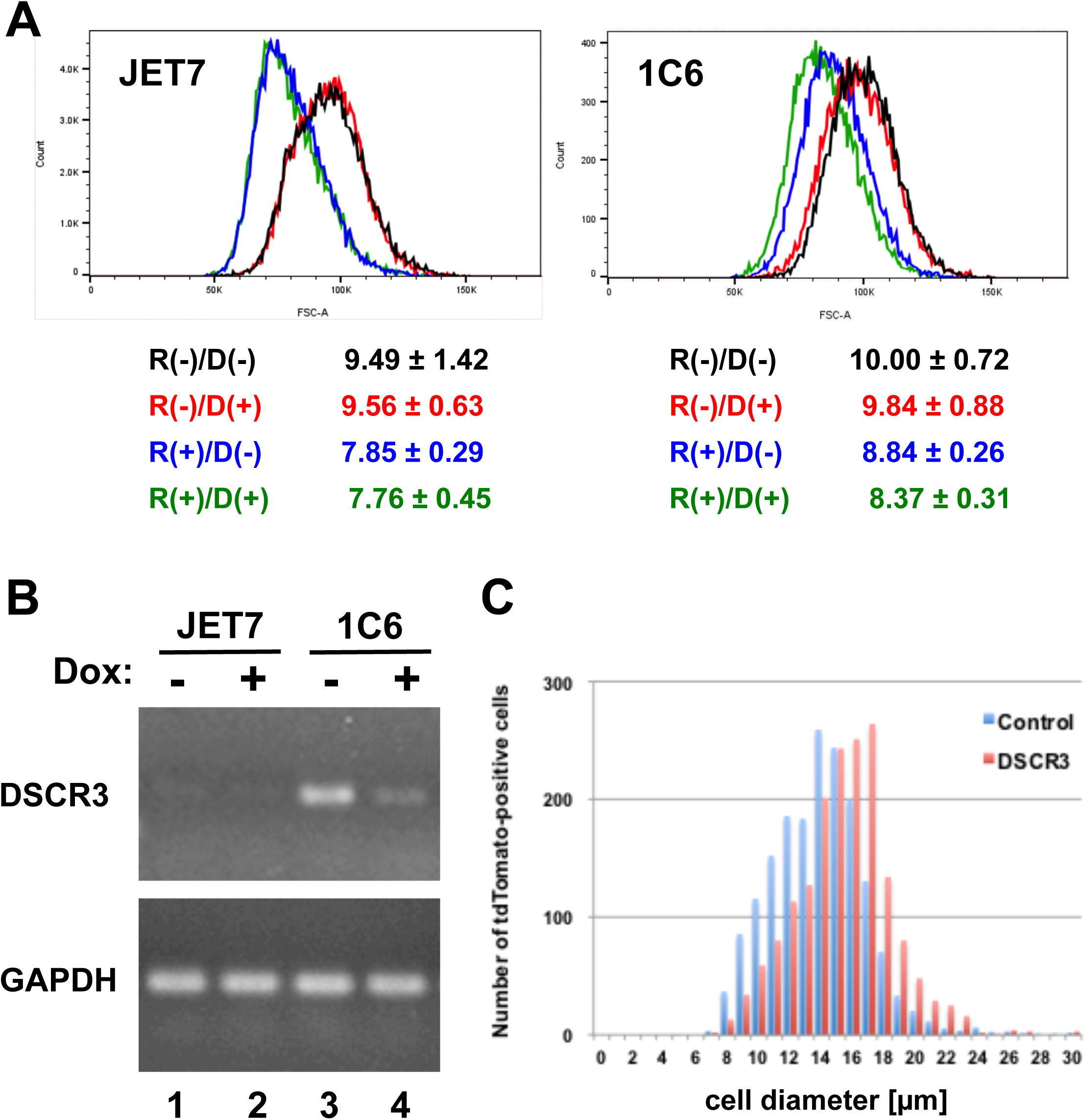
Isolation and characterization of DSCR3 as a cell size-controlling gene. (A) Cell size distributions of control JET7 cells (left panel) and JET7-derived clone 1C6 cells (right panel). Both cell types were untreated [R(-)/D(-), black lines], treated with doxycycline alone [R(-)/D(+), red lines], treated with rapamycin alone [R(+)/D(-), blue lines], or treated with both [R(+)/D(+), green lines], and the cell size distribution was measured as forward scatter (FSC) by flow cytometry. One representative result is shown for the four independent experiments. An example of the gating strategy for doublet exclusion is shown in Supplementary figure 1. The mean FSC values for each treatment are indicated below the panels. Standard deviations were calculated from three consecutive measurements. (B) Total RNA was extracted from JET7 and 1C6 cells in the presence or absence of doxycycline (indicated by Dox: + or -), and the expression level of DSCR3 was examined using reverse transcription-PCR (upper panel). GAPDH levels were measured as a control (lower panel). (C) HEK293 cells were transfected with a DSCR3-overexpressing plasmid (DSCR3) as well as the control empty plasmid (Control), and the cell size distribution for the individual transfectants is indicated (Control, blue bars; DSCR3, red bars). One representative data from four independent transfection experiments is shown.

We then analyzed the ERM-trapped transcript and identified the Down syndrome critical gene 3 (DSCR3) as the gene responsible for clone 1C6. We observed qualitative upregulation of DSCR3 transcription in 1C6 cells in a tTA-dependent manner (Figure 1B). When we introduced a DSCR3-overespression plasmid into HEK293 cells, the size distribution shifted to a larger area than that observed in cells transfected with the control empty plasmid (Figure 1C). To evaluate the difference in cell size between the Control and the DSCR3-overexpressing cells, we conducted Welch’s *t*-test using the mean cell size obtained from four independent transfection experiments. The mean cell size was 14.71 (SD = 0.76) for Control and 15.84 (SD = 0.48) for DSCR3, which did not reach statistical significance at the conventional 5% level (*p* = 0.0525). However, the analysis demonstrated a very large effect size (Hedges’ *g* = 1.55), indicating that practically substantial differences in cell size existed between the two groups. Thus, we concluded that exogenous overexpression of DSCR3 induced cell enlargement.

### Contribution of DSCR3 for cell size of the fibroblasts from individuals with DS

DSCR3 is a gene located on human chromosome 21q22 that is critical for the characteristic features of DS. Therefore, we examined the mRNA levels of DSCR3 in fibroblasts derived from individuals with DS. The expression of DSCR3 relative to that of GAPDH was approximately 33% higher in fibroblasts from individuals with DS (Figure 2A, Detroit 529 and 532) than in fibroblasts obtained from healthy donors (Figure 2A, TIG-119 and 120). Indeed, the protein level of DSCR3 was elevated in DS fibroblasts compared to that in normal fibroblasts (Figures 2B and 2C). Interestingly, immunofluorescence images of fibroblasts revealed that DS fibroblasts were larger than normal ones (Figure 2D). Cells were detached from the culture plates, and their electronic cell volume (EV) was measured using a Coulter counter. Although TIG-120 did not show significant differences in EV compared to any other cell line, the DS fibroblasts were statistically larger than TIG-119. These results indicate that DSCR3 was overexpressed in DS fibroblasts, contributing to the increase in cell volume.

**FIGURE 2.**
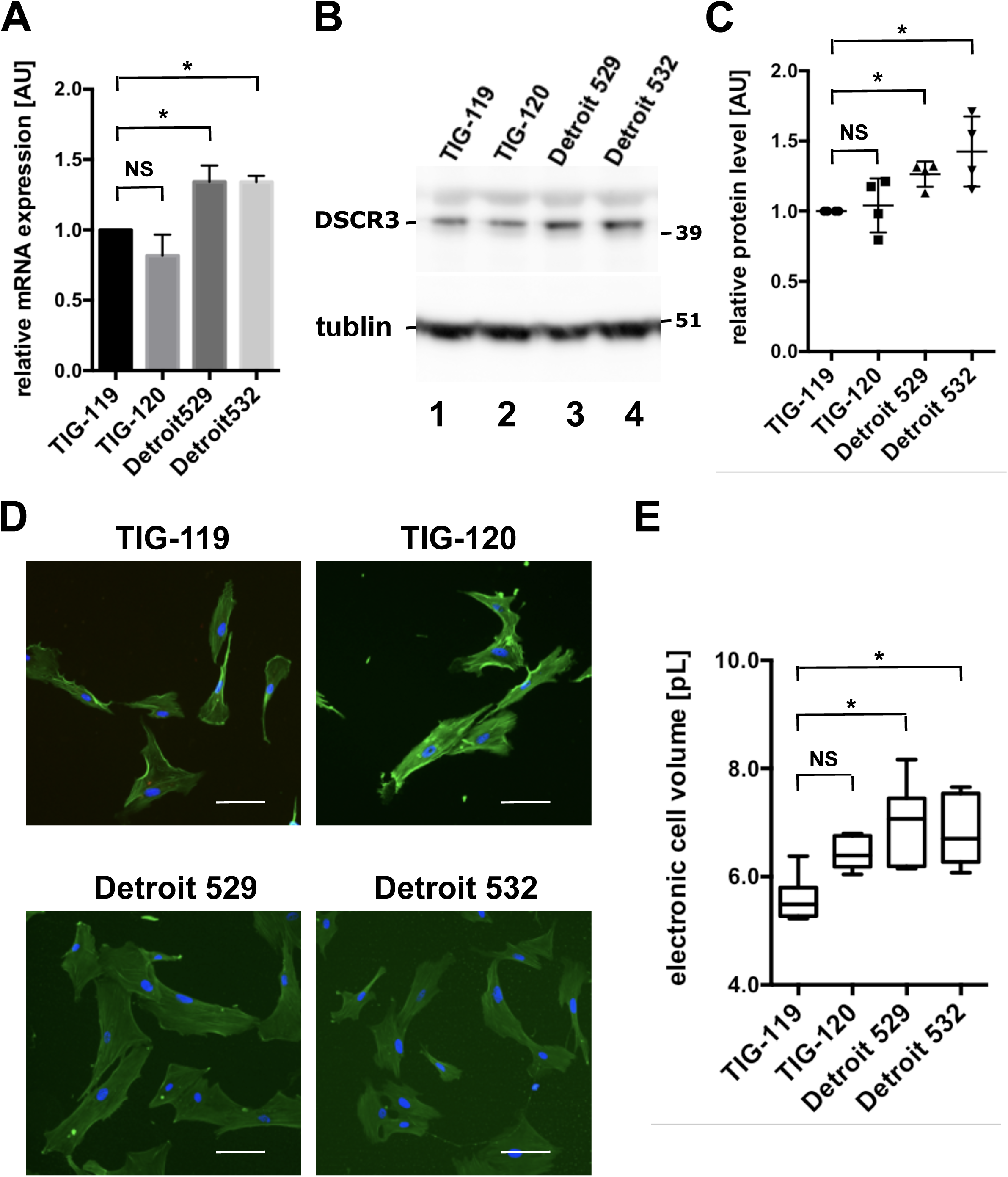
Characterization of DSCR3 in fibroblasts from normal and DS individuals. (A) Relative expression of DSCR3 mRNA to GAPDH mRNA was quantified in fibroblasts from healthy donors (TIG-119 and TIG-120) and individuals with DS (Detroit 529 and Detroit 532). The bar graph represents the averages of three experiments with standard deviations. Statistical analysis was performed using One-way ANOVA followed by Tukey’s HSD post-hoc test for multiple comparisons. NS, not significant (*p* > 0.1); *, *p* = 0.011. (B) Immunological detection of DSCR3 protein in fibroblasts from healthy donors (TIG-119 and TIG-120) and individuals with DS (Detroit 529 and Detroit 532). Tubulin was detected using the same blot (lower panel). The position of the cognate protein is shown on the left and that of the molecular weight marker is shown on the right of the blot. One representative result is shown for the four independent experiments. (C) Relative protein levels of DSCR3 to α-tubulin were calculated and plotted from four independent experiments. Error bars represent the standard deviation of four independent experiments. NS, not significant (*p* > 0.5); *, *p* < 0.02, calculated by unpaired *t*-test. (D) Fluorescence images of fibroblasts from healthy donors (TIG-119 and TIG-120) and individuals with DS (Detroit 529 and Detroit 532) stained with phalloidin for F-actin (green) and DAPI for nuclei (blue). The white horizontal bar in each image represents 100 µm. (E) Electronic cell volumes of fibroblasts from healthy donors (TIG-119 and TIG-120) and individuals with DS (Detroit 529 and Detroit 532) were measured using a Coulter Counter. Average cell volumes measured across six independent experiments are presented in boxplots with standard deviations. A One-way ANOVA was conducted to test for overall differences among the cell lines, followed by Tukey’s HSD post-hoc test for multiple comparisons. NS, not significant (*p* > 0.05); *, *p* < 0.01.

### Impact of DSCR3 for cell growth, respiration, and size control

Next, we examined the growth of DS and normal fibroblasts. Although the changes in cell number were comparable at early time points (Figure 3A, 30 and 66 hours), differences in the growth rate between DS and normal fibroblasts became evident at later stages (Figure 3A, 102 and 138 hours). As shown in Supplementary Figure 2A, Detroit 529 cells were still subconfluent even at 138 hours, indicating that the lower growth rates of DS fibroblasts were not caused by contact inhibition. In fact, doubling times of DS fibroblasts (43.67 ± 2.70 hours for Detroit 529 and 54.91 ± 2.10 hours for Detroit 532) were statistically longer than those of normal fibroblasts (32.05 ± 2.80 hours for TIG-119 and 36.72 ± 1.50 hours for TIG-120). Mitochondrial respiration was compared using an extracellular flux analyzer. In contrast to cell growth capacity, both basic and maximal respiration were higher in DS fibroblasts than in normal ones (Figure 3B and Supplementary figure 2). Finally, we investigated the effects of silencing endogenous DSCR3 using siRNA-mediated knockdown. We used a mixture of four independent siRNAs targeting DSCR3 (siPOOL), one specific siRNA (siINDI), and a non-targeting control siRNA (siCONT) for the transfection. DS and normal fibroblasts were transfected with a set of siRNAs, and the cell size was measured. The cell sizes of siCONT-transfected cells were reduced compared to those of non-transfected cells (c.f. Figure 2E). This may be caused by the transfection reagent rather than the off-target effects of siRNA, since we reproducibly observed such cell size reductions throughout the transfection experiments, even in the absence of siRNA (data not shown). Knockdown of DSCR3 by siPOOL and siINDI effectively reduced the protein level of DSCR3 to 51 % of that induced by siCONT (Supplementary figure 3), as well as the cell size in both DS and normal fibroblasts. However, we could not show a significant difference in cell size for Detroit 532, probably due to the scattered distribution of cell size in the control siRNA-transfected cells (Figure 3C). The degree of cell size reduction, calculated as the %-reduction rate of cell volume against individual control siRNA-transfected cells, was not significantly different among the cell lines tested (*p* value of one-way ANOVA = 0.7169; Figure 3D). These results suggest that DSCR3 contributes equally to the regulation of cell size in both DS and normal fibroblasts.

**FIGURE 3.**
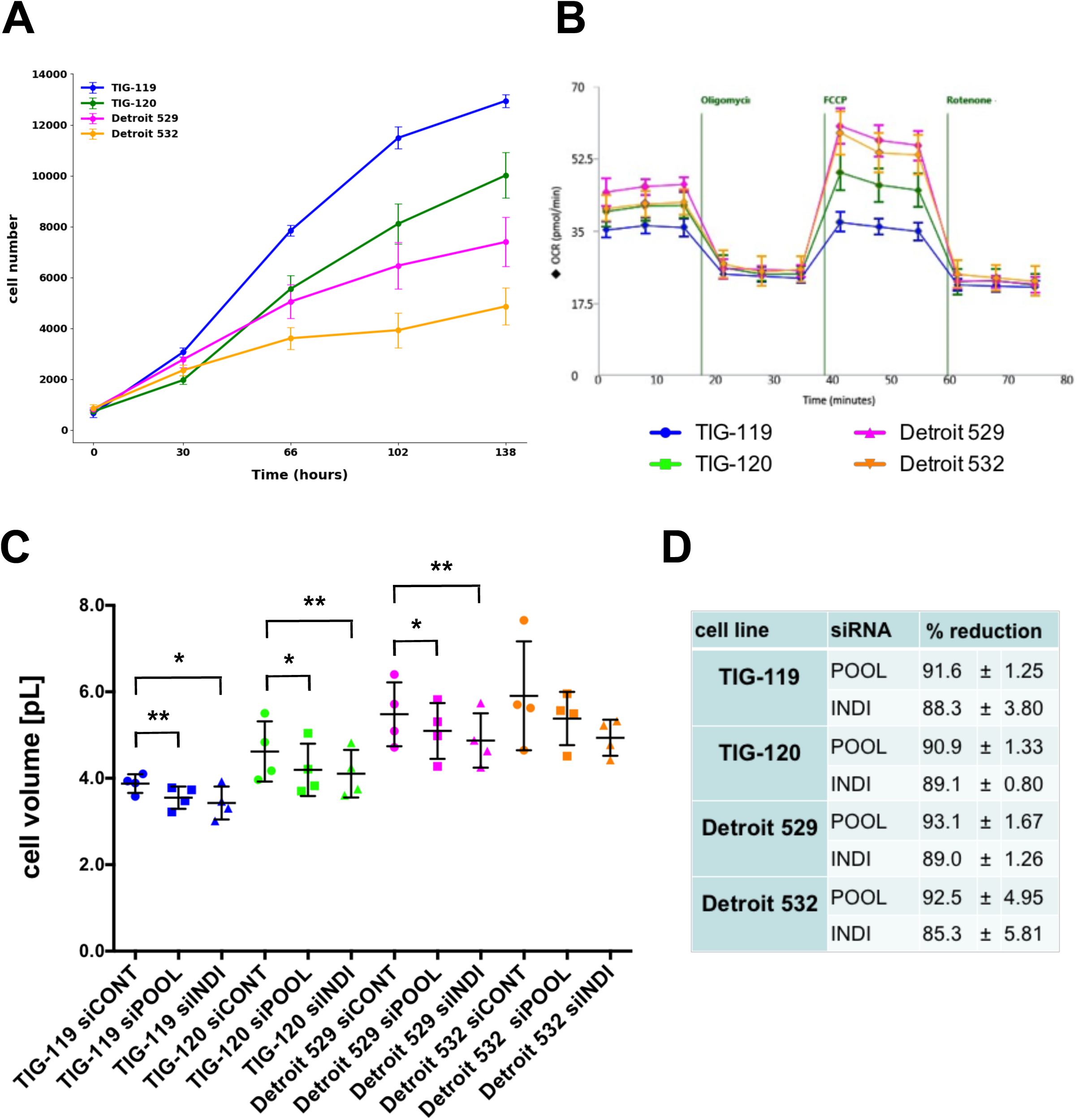
Characterization of fibroblasts from normal and DS individuals. (A) Comparison of the cell numbers for each cell line at 30, 66, 102, and 138 hours from the beginning (0 h) of the observation. Each spot represents the average cell number from five wells in a 96-well plate for each cell line, with standard deviation. (B) Mitostress assay was performed on fibroblasts from healthy donors (TIG-119 and TIG-120) and DS individuals (Detroit 529 and Detroit 532) using an extracellular flux analyzer. The averages from the eight wells were plotted with standard deviations. (C) Cell volume measurements of fibroblasts from healthy donors (TIG-119 and TIG-120) and DS individuals (Detroit 529 and Detroit 532) transfected with control (siCONT) or DSCR3-targeting siRNAs (siPOOL and siINDI). Paired *t*-tests were performed for individual cell lines between DSCR3-targeting siRNA transfectants and control siRNA transfectants. *, *p* < 0.055; **, *p* < 0.01. (D) Cell size reduction of fibroblasts from healthy donors (TIG-119, TIG-120) and DS individuals (Detroit 529, Detroit 532) treated with DSCR3-targeting siRNAs (siPOOL, siINDI) compared to that of control siRNA. The data are presented as the average %-reduction in the electronic cell volume of DSCR3-targeting siRNA-treated cells compared to that of the control siRNA-treated cells from four independent transfection experiments.

## Discussion

Here, we describe DSCR3, which functions as a cell size-controlling gene, identified using a novel cell size mutant screening strategy. Overexpression of DSCR3 enlarged cells using both the ERM-tagged system and transient transfection methods. Rapamycin treatment reduced the size of both control and DSCR3-overexpressing cells, but the size distribution of the latter remained larger than that of the former. However, the reduction rate was similar for both cell types, suggesting that cell size regulation by DSCR3 is independent of the mTOR signaling pathway. Consistent with this notion, siRNA-mediated knockdown of DSCR3 resulted in the same degree of reduction in cell size in both normal and DSCR-overexpressing fibroblasts from individuals with DS, although it does not exclude the possibility that factors other than DSCR3, particularly one(s) in the DSCR, contribute to cell size enlargement in DS fibroblasts. In addition, it may be interesting to investigate how cell size reduction affects cell growth, mitochondrial respiration, and protein transport in normal and DS fibroblasts.

The DS critical region (DSCR) is a segment of chromosome 21 hypothesized to contain genes responsible for many features of DS, including cognitive impairment and craniofacial abnormalities (Dierssen, 2012). The DSCR contains approximately 20 genes, including RCAN1 (Regulator of Calcineurin 1) as DSCR1 and PSMG1 (proteasome assembly chaperone 1) as DSCR2. Although the roles of RCAN1/DSCR1 in multiple signaling pathways and their possible contribution to the DS phenotype have been extensively discussed (Lee and Ahnn, 2020), the role of PSMG1/DSCR2 remains poorly understood. At the time we determined that DSCR3 functions as a cell size regulator, no biological functions were reported in the common database. A later study revealed that DSCR3 is a component of the cargo complex that plays a crucial role in endosomal cargo retrieval and membrane protein recycling (Boesch et al., 2024; Mcnally et al., 2017). The retriever complex is a heterotrimer composed of DSCR3, C16orf62, and VPS29, which shows a striking structural similarity to another cargo protein complex, the retromer, which recycles transmembrane receptors from endosomes to the trans-Golgi network. Therefore, it has been proposed that DSCR3 and C16orf62 be renamed VPS26C and VPS35L, respectively (McNally et al., 2017). Although the functional relevance of VPS26C/DSCR3 to the DS phenotype remains to be investigated, the possible link between endosomal protein recycling and cell size regulation is intriguing.

It is also important to note that the growth rate of DS fibroblasts is slower than that of normal fibroblasts (Figure 3A; Kawakubo et al., 2017), whereas the mitochondrial respiration capacity is higher in the former. Indeed, fibroblasts differentiated from DS individual-derived induced pluripotent stem cells also showed higher mitochondrial activity (Nawa et al., 2019; Adorno et al., 2013). As the size of human fibroblasts increases, their proliferative capacity decreases (Neurohr et al., 2019). Therefore, one possible reason for the slow proliferation of fibroblasts derived from DS individuals may be their increased size. Increased cell size dilutes cytoplasmic protein concentrations (Neurohr et al., 2019) and induces aging-like proteome changes in non-aging cells (Lanz et al., 2022), leading to cellular senescence (Chadha et al., 2024). As DSCR3 is expressed in tissues throughout the body, it may promote systemic aging in individuals with DS by causing cell enlargement throughout the body. In fact, DS is considered a form of premature aging (Horvath et al., 2015). Although premature aging in individuals with DS is unlikely to be explained by a single gene, DSCR3 is an optimal target for inhibiting premature aging in individuals with DS. Additionally, basal and maximal respiration were higher in fibroblasts derived from individuals with DS than in those derived from healthy controls (Figure 3B). The mitochondrial membrane potential varies with cell size and is independent of the cell cycle (Miettinen and Björklund, 2016). Therefore, the increased basal and maximal respiration in fibroblasts derived from individuals with DS may be correlated with their cell size. However, since cellular aging leads to a decrease in respiratory levels and an increase in reactive oxygen species stress (Hutter et al., 2004), the peculiar relationship between cell size expansion, increased respiratory levels, and cellular aging in DS fibroblasts requires further investigation.

### Experimental Procedures

#### Cell culture

A Jurkat-derived cell line stably expressing the tetracycline-responsive transactivator tTA, JET7, and the cell size-mutant clone 1C6 were cultured in RPMI1630 (Wako Pure Chemical Industries, Ltd., Osaka, Japan) containing 10% fetal bovine serum (FBS; Peak Serum, Inc., Bradenton, FL, U.S.A.), 100 U/mL penicillin/100 µg/mL streptomycin (Wako Pure Chemical Industries, Ltd., Osaka, Japan) in the presence of 1 mg/ml Geneticin (Sigma-Aldrich Co., LLC, St. Louis, MO, U.S.A.) and 0.2 mg/ml Hygromycin B (Nacalai Tesque, Inc., Kyoto, Japan). HEK293 cells were cultured in Dulbecco’s Modified Eagle Medium (D-MEM; Nacalai Tesque, Inc., Kyoto, Jpan) containing 10% FBS (Peak Serum, Inc., Bradenton, FL, U.S.A.), 100 U/mL penicillin/100 µg/mL streptomycin (Wako Pure Chemical Industries, Ltd., Osaka, Japan). TIG-119 and TIG-120 derived from 6-month-old male and female Japanese healthy controls, respectively, and Detroit 532 derived from 2-month-old male Caucasian children with DS were obtained from the Japanese Collection of Research Bioresources (JCRB) Cell Bank (Osaka, Japan) (Kawakubo et al., 2017). Detroit 529 cells derived from 2.5-year-old female Caucasian children with DS were obtained from (American Type Culture Collection) (Manassas, VA, U.S.A.). TIG-119 and TIG-120 cells were cultured in Eagle’s minimum essential medium (EMEM; Wako Pure Chemical Industries, Ltd., Osaka, Japan) containing 10% fetal bovine serum (FBS; SAFC Biosciences, Inc., Lenexa, KS, U.S.A.) and 100 U/mL penicillin/100 µg/mL streptomycin (Nacalai Tesque, Inc., Kyoto, Japan). Detroit 529 and Detroit 532 cells were cultured in EMEM containing 10% FBS, 100 U/mL penicillin/100 µg/mL streptomycin, nonessential amino acids (Nacalai Tesque, Inc.), 1 mM sodium pyruvate (Nacalai Tesque, Inc., Kyoto, Japan), and 0.1% lactalbumin hydrolysate (Sigma-Aldrich Co., LLC, St. Louis, MO, U.S.A.). All cells were cultured at 37°C in a humidified atmosphere containing 5% CO_2_.

#### Cell size measurements

JET7 and 1C6 cells were cultured as described in the presence and/or absence of 20 nM rapamycin (Cell Signaling Technology, Danvers, MA, U.S.A.) and 1 µg/mL doxycycline (Sigma-Aldrich Co., LLC, St. Louis, MO, U.S.A.), as described in the figures. Cells were analyzed by flow cytometry using FACS Canto II (Becton, Dickinson and Company, Franklin Lakes, NJ, U.S.A.), and their size distributions were characterized as forward scatter. Cell size distribution was measured using a CellDrop FL automated cell counter (SCRUM Inc., Tokyo, Japan) or a Moxi-Z cell counter (ORFLO Technologies, Livermore, CA, U.S.A.) according to the manufacturer’s instructions.

#### RNA isolation and RT-PCRs

For quantitative reverse transcription-PCR, total RNA was purified from JET7 and 1C6 cells using the RNeasy Mini Kit (QIAGEN, Venlo, The Netherlands), and 100 ng of total RNA was reverse-transcribed using the StrataScript First-Strand Synthesis Kit (Stratagene, La Jolla, CA, U.S.A.). PCR amplification was performed using HotStarTaq DNA polymerase (QIAGEN, Venlo, The Netherlands). Thermocycling (95°C for 15 min, followed by 30 cycles of 95°C for 1 min, 58°C for 1 min, and 72°C for 1 min) was performed using a Mastercycler (Eppendorf, Hamburg, Germany). For quantitative real-time PCR, total RNA was purified from human fibroblasts using the High Pure RNA Isolation Kit (Roche Diagnostics, Mannheim, Germany), and 800 ng of total RNA was reverse transcribed using PrimeScript RT Master Mix (Takara Bio, Shiga, Japan) and assessed by quantitative real-time PCR using tubulin mRNA as an internal control. PCR amplification was performed using SYBR Premix Ex Taq (Tli RNaseH Plus) (Takara Bio, Shiga, Japan). Thermocycling (95°C for 30 s, followed by 40 cycles of 95°C for 5 s and 60°C for 30 s) was performed using an Applied Biosystems 7300 Real-Time PCR System (Applied Biosystems) (Waltham, MA, U.S.A.). The following primers were used for quantitative real-time PCR amplifications: *DSCR3*: forward: 5’-AAGGAGGGAGGTGTGTGAGGTGT-3’, reverse: 5’-TGAAGAGCAGATGACAAACGCCC-3’; *GAPDH*: forward: 5’-GCACCGTCAAGGCTGAGAAC-3’, reverse: 5’-TGGTGAAGACGCCAGTGGA-3.’

#### Construction of the plasmid and transfection

The construction of the plasmid overexpressing DSCR3 has been described in the literature (Kato et al., in press). Transfection of the plasmid (DSCR3_ptdTomato) and control vector (Δ NN_ptdTomato) was performed using Lipofectamine 3000 in Opti-MEM, according to the manufacturer’s protocol. The cell sizes of tdTomato-positive cells were measured as described above.

#### Western blotting

Human fibroblasts were cultured in 10 cm plates at subconfluence and recovered in PBS(-) using a cell scraper. Cells were collected by centrifugation and lysed in a buffer containing 40 mM HEPES (pH 7.5), 120 mM NaCl, 1 mM EDTA, 0.3% CHAPS, supplemented with PhosSTOP, and cOmplete Protease Inhibitor Cocktail without EDTA (Sigma-Aldrich Co., LLC, St. Louis, MO, U.S.A.). After gentle shaking at 4°C for 10 min, the cell suspensions were centrifuged at 100,000 × g for 5 min at 4°C. The supernatants were recovered and applied onto an SDS-PAGE gel, followed by western blotting using an XCell SureLock mini cell electrophoresis system (ThermoFisher Scientific, Waltham, MA, U.S.A.). Anti-DSCR3 polyclonal antibody (#AP6319a; Abgent, San Diego, CA, U.S.A.) and anti-α-tubulin monoclonal antibody (clone DM1A, #3873; Cell Signaling Technology, Danvers, MA, U.S.A.) were used as primary antibodies.

#### Immunofluorescence

Human fibroblasts were cultured in collagen-coated glass-bottom chamber slides at 2×10^4^ cells/area overnight. The cells were fixed with 4% paraformaldehyde. After washing with PBS(-), the cells were permeabilized with 0.5% Triton X-100 in PBS(-) for 5 min at room temperature. Cells were washed and blocked with 1% BSA recovered in PBS(-) for 15 min at RT. The blocking solution was then replaced with a primary antibody diluent and incubated for 1 h at room temperature. After washing with PBS(-), the cells were incubated with a fluorescently labeled secondary antibody diluent for 30 min at room temperature. The cells were washed and successively soaked in 70%, 80%, 90%, and 100% ethanol. After drying the slides, the cells were covered with slide glass and VECTASHIELD mounting medium. Fluorescent images were obtained using a Keyence BZ-X700 fluorescent microscope (Keyence Corporation, Osaka, Japan).

#### Cell growth and flux analysis

A 96-well plate containing human fibroblasts at 1×10^3^ cells/well was placed in a live cell imager (IncuCyte ZOOM; Sartorius, Göttingen, Germany) in a humidified CO_2_ incubator at 37°C. Time-lapse images were obtained every 6 h for 3 days. After changing the culture medium, the cells were incubated and monitored for two days. The number of cells in each image was counted in the presence of IncuCyte Cytotox Red reagent (Sartorius, Göttingen, Germany) and plotted according to the manufacturer’s instructions. For extracellular flux measurements, cells were seeded at 2.8×10^4^ cells/well in an assay plate and incubated overnight in a humidified CO_2_ incubator at 37°C. After replacing the culture medium with serum-free assay medium, the plate was set in an extracellular flux analyzer XFe96 (Seahorse Biosciences, North Billerica, MA, U.S.A.) and applied for the mitostress assay with 1 µM oligomycin, 0.5-1.0 µM FCCP, and 0.125 µM rotenone and antimycin.

#### Transfection of siRNA

A total of 80,000 fibroblasts were seeded in a 6-well plate and cultured overnight. After replacing the culture medium with an antibiotic-free medium, the siRNAs were transfected using Lipofectamine RNAiMAX (Thermo Fisher Scientific, Waltham, MA, U.S.A.) according to the manufacturer’s instructions. The following siRNAs were used: ON-TARGETplus Human DSCR3 (10311) siRNA SMARTpool (L-012163-02-0005) and Individual (J-012163-19-0002), and Non-targeting siRNA Control#2 (D-001810-02-05) (Horizon Discovery, Cambridge, U.K.). The electronic volume of the transfected cells was measured three days after transfection, as described above.

#### Statistical Analysis

Statistical analysis was performed as described in the figure legends using Microsoft Excel or GraphPad PRISM. Unless otherwise indicated, all values are expressed as mean ± standard deviation. The significance thresholds are indicated in the figure legends and text.

## Author Contributions

**Kazumi Kimura:** formal analysis, investigation. **Masakazu Souda:** data curation, investigation. **Ryotaro Mori:** investigation. **Yoshimi Kato:** materials. Hiroki Kurahashi: materials. **Masashi Asai:** writing – review and editing. **Kazuo Yamamoto:** conceptualization, data curation, formal analysis, funding acquisition, investigation, methodology, project administration, resources, validation, visualization, writing – original draft preparation, writing – review and editing.

## Acknowledgements

We thank Dr. Katsuya Hirasaka for the use of an equipment. Ms. R. Miyazaki, Y. Iwasaki, Y. Yamasaki and K. Saiki for their support.

## Ethics Statement

The authors have nothing to report.

## Data Availability Statement

The data supporting the findings of this study are available from the corresponding author upon reasonable request.

