## Supplementary Figures for "Identification of the Down syndrome critical region 3 gene as a mammalian cell size regulator"

### Supplementary figure 1

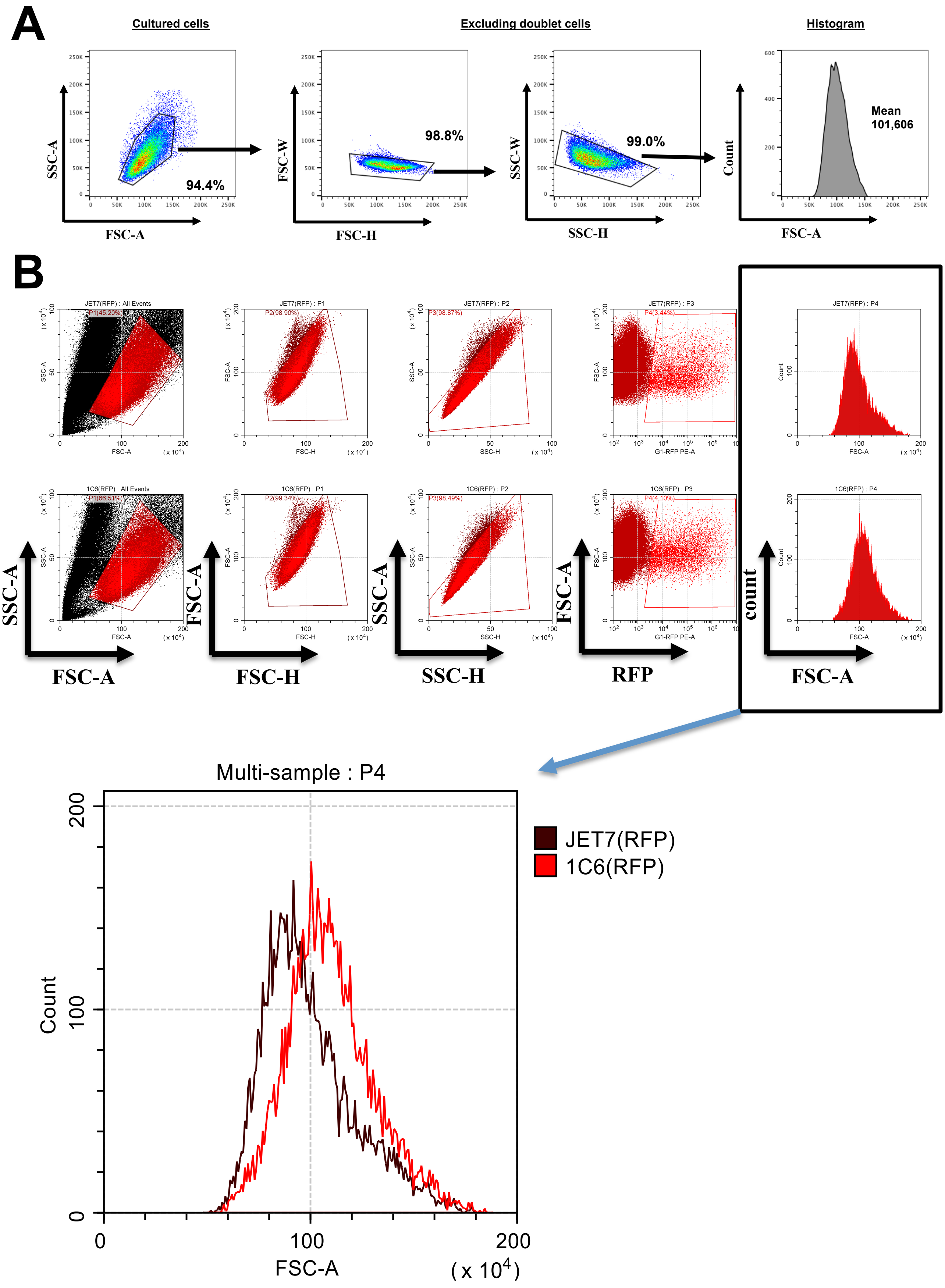

### Supplementary figure 2

A

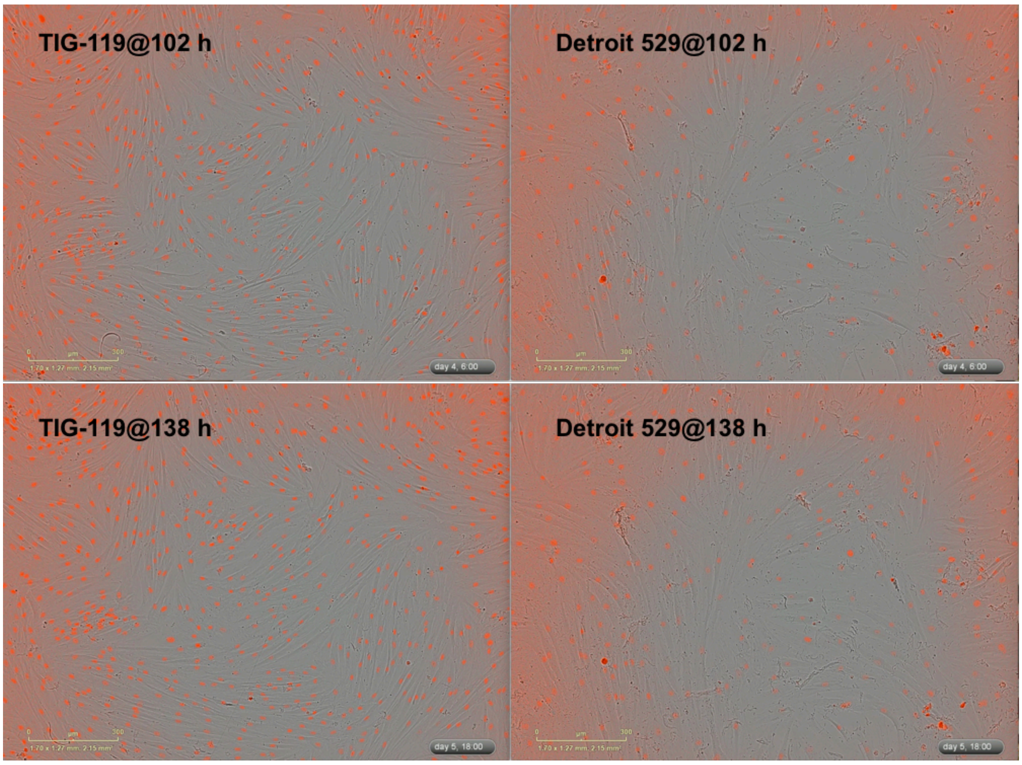


B

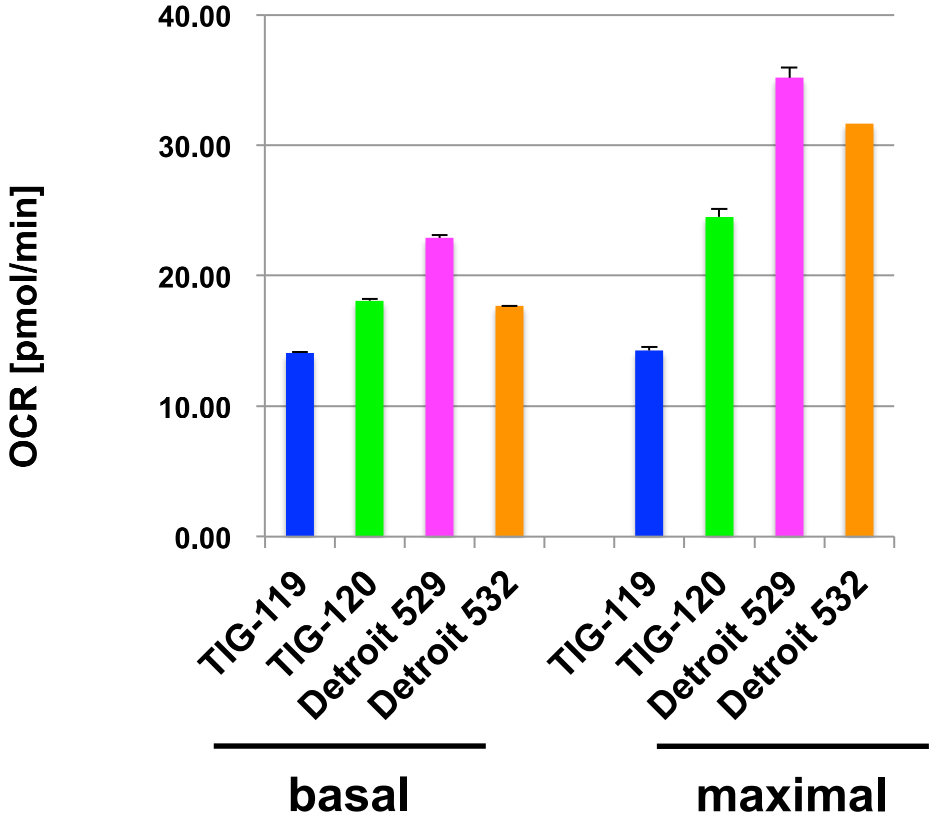

### Supplementary figure 3

| Cell line | siRNA | Relative protein level [%] | Average | Deviation |
|---|---|---|---|---|
| TIG119 | control | 100.0 | 100.0 | 0.0 |
| TIG120 | control | 100.0 | | |
| Detroit529 | control | 100.0 | | |
| Detroit532 | control | 100.0 | | |
| TIG119 | DSCR3 pool | 52.3 | 50.9 | 7.2 |
| TIG120 | DSCR3 pool | 50.1 | | |
| Detroit529 | DSCR3 pool | 37.1 | | |
| Detroit532 | DSCR3 pool | 63.9 | | |
| TIG119 | DSCR3 individual | 65.8 | 51.2 | 8.0 |
| TIG120 | DSCR3 individual | 36.0 | | |
| Detroit529 | DSCR3 individual | 50.4 | | |
| Detroit532 | DSCR3 individual | 52.5 | | |

**Supplementary figure legends**

**Supplementary figure 1** (A) Example of the gating strategy for doublet exclusion. JET7 treated with both rapamycin and doxycycline alone [R(+)/D(+), green line in the left panel, FIGURE 1A] is shown. (B) Cell size distribution of JET7 and 1C6 cells in the G1 phase. Both cell lines were transfected with a plasmid expressing mCherry-Ctd1 to label the G1-cell populations. The gating strategy and resultant cell size distributions are shown.

**Supplementary figure 2** (A) Representative images of Detroit 529 at 102 and 138 hours from the beginning (0 h) of the observation. Reddish backgrounds originated from IncuCyte Cytotox Red reagent to spot live cells, which did not interfere with cell growth and viability. (B) Oxygen Consumption Rates (OCR) for basal and maximal respiration are summarized as a bar graph representation. Data from the first three points (before adding oligomycin) or those obtained between the addition of FCCP and the addition of rotenone/antimycin B were averaged and indicated with standard deviation.

**Supplementary figure 3** Summary of relative DSCR3 protein levels in individual siRNA-treated fibroblasts. The signals of DSCR3 in fibroblasts from normal and DS individuals were quantified and normalized against those of $\alpha$-tubulin. The data averaged from four immunoblots were compared between siCONT- and siPOOL-treated cells or between siCONT- and siINDI-treated cells, respectively (= Relative protein level [%]). The data of for individual siRNA-treated cells were further averaged (= Average) to compare the impact of individual siRNAs on cell size reduction (Deviation = standard deviation).
